# Extracting Brain Disease-Related Connectome Subgraphs by Adaptive Dense Subgraph Discovery

**DOI:** 10.1101/2020.10.07.330027

**Authors:** Qiong Wu, Xiaoqi Huang, Adam Culbreth, James Waltz, Elliot Hong, Shuo Chen

**Affiliations:** Department of Mathematics, University of Maryland, College Park, MD, USA; Department of Mathematics, Johns Hopkins University, Baltimore, MD, USA; Maryland Psychiatric Research Center, School of Medicine, University of Maryland, Baltimore, MD, USA

**Keywords:** brain connectome, densest subgraph, likelihood-based criterion, network topology, permutation test, schizophrenia

## Abstract

Group-level brain connectome analysis has attracted increasing interest in neuropsychiatric research with the goal of identifying connectomic subnetworks (subgraphs) that are systematically associated with brain disorders. However, extracting disease-related subnetworks from the whole brain connectome has been challenging, because no prior knowledge is available regarding the sizes and locations of the subnetworks. In addition, neuroimaging data is often mixed with substantial noise that can further obscure informative subnetwork detection. We propose a likelihood-based adaptive dense subgraph discovery (ADSD) model to extract disease-related subgraphs from the group-level whole brain connectome data. Our method is robust to both false positive and false negative errors of edge-wise inference and thus can lead to a more accurate discovery of latent disease-related connectomic subnetworks. We develop computationally efficient algorithms to implement the novel ADSD objective function and derive theoretical results to guarantee the convergence properties. We apply the proposed approach to a brain fMRI study for schizophrenia research and identify well-organized and biologically meaningful subnetworks that exhibit schizophrenia-related salience network centered connectivity abnormality. Analysis of synthetic data also demonstrates the superior performance of the ADSD method for latent subnetwork detection in comparison with existing methods in various settings.

## 1 Introduction

Brain connectome analysis has become a powerful tool to understand the neurophysiology and neuropathology of brain diseases at a circuit level. These analyses focused on investigating patterns of functional and/or structural inter-connections between neural populations in the central nervous system associated with symptomatic phenotypes. Mounting evidence has shown that major neuropsychiatric disorders, including schizophrenia, Alzheimer’s disease, and autism among others, are associated with disrupted structural and functional connectivity patterns (Fornito et al., 2012).

Recent advances in neuroimaging statistics have facilitated group-level statistical analysis of structural and functional brain connectome data and the identification of disease-related brain connectome patterns (Biswal et al., 2010, Cao et al., 2019, Chen et al., 2019). In these analyses, the brain is often depicted as a graph (Bullmore and Bassett, 2011), where each node corresponds to a brain region of interest (ROI) and an edge represents the connectivity linking any two nodes. An edge can represent functional connectivity based on functional magnetic resonance imaging (fMRI) data at rest or task, structural connectivity measuring white matter track connections, and weighted connection metric integrating multimodal brain connectivity (Bowman et al., 2012). These multivariate edges are the variables of interest in brain connectome analysis, which are constrained by the nodes in a weighted adjacency matrix and thus exhibit network topological properties (Chen et al., 2020). Statistical inference for multivariate edge variables in an adjacency matrix remains challenging because of the need to account for multiple testing corrections and network topological structures simultaneously. Many statistical graph models have been developed and successfully applied to brain connectome data analysis yielding important findings (Vogelstein et al., 2012, Durante et al., 2018, Ghoshdastidar and von Luxburg, 2018, Ginestet et al., 2017, Higgins et al., 2019, Kundu et al., 2018, Lukemire et al., 2017, Mejia et al., 2019, Simpson et al., 2019, Xia and Li, 2017, Xia and Li, 2018 among many).

The current study focuses on extracting informative/signal subgraphs that are likely related to brain diseases from the whole brain connectome (Vogelstein et al., 2012). Our overarching goal is to accurately capture underlying signal subnetworks, such that the extracted subnetworks i) cover a high proportion of true positive edges (i.e., high sensitivity); (ii) include a few false positive edges (i.e., low false discovery rate (FDR)); and iii) are composed of highly organized network topological structures (i.e., biologically interpretable). In practice, however, this task is challenging because it is difficult to simultaneously balance the sensitivity and false positive findings while constraining all positive edges in organized subgraphs. The ‘dense’ subnetwork detection then becomes attractive because a subgraph of a small number of nodes in an organized network topological structure covering most signal edges can also lead to low FDR and high sensitivity. Although less discussed in the statistical literature, dense subgraph discovery in the field of computer science research has been carefully worked out (Lee et al., 2010), and thus may suit our needs for statistical analysis of brain connectome data.

Dense subgraph discovery methods are designed to identify a subgraph with a maximal density among all possible subgraphs, in short, the densest subgraph, in a binary graph. These methods rely on the assumption that the overall graph is non-random and there exists some subgraph where the edge ratios are much higher than the rest of the graph. Goldberg (1984) reduces the problem to a sequence of max-flow/min-cut computations, which requires a logarithmic number of min-cut calls. Asahiro et al. (1996) propose a simple and fast greedy algorithm that is showed to have a 2-approximation guarantee by Charikar (2000). Nevertheless, existing dense subgraph discovery algorithms may not be directly applicable to our analysis due to the substantial noise in the group level brain connectome data. As demonstrated in Figure 1a, there may exist an enormous amount of false positive and false negative errors in edge-wise inference results that give rise to the difficulty of detecting dense subgraph using existing methods. Specifically, due to the noise, existing dense subgraph discovery algorithms tend to either identify over-sized subgraphs that may include a large proportion of false positive edges with low importance levels (high FDR) or detect over-conservative small-sized subgraphs that may not sufficiently cover signal edges (low sensitivity, Tsourakakis et al., 2013). Moreover, the computational cost of many these dense subgraph density discovery algorithms is expensive, which may lead to intractable computational time for commonly used statistical inference methods for brain connectome analysis (e.g., permutation tests). Hence, we are motivated to integrate modern statistical techniques into dense graph discovery and mitigate these challenges for brain connectivity network analysis.

**Figure 1:**
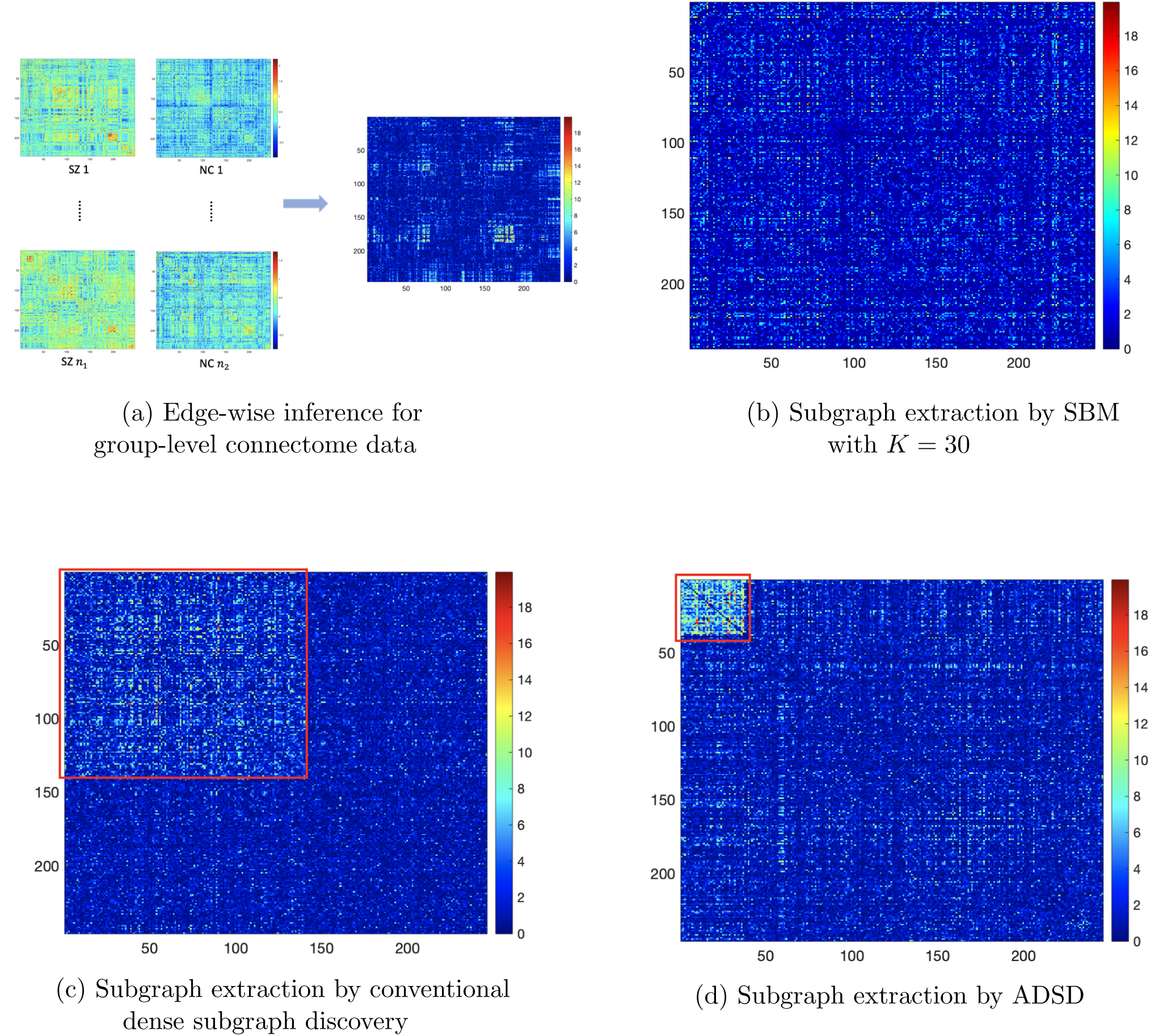
Motivation for informative subgraph extraction: (a) demonstrates the process of obtaining edge-wise inference matrix from the population level connectome data; (b) illustrates the commonly used community detection results (e.g. using stochastic block model) cannot detect any informative subgraph; (c) shows the results of existing dense subgraph discovery results; (d) describes a desirable informative subgraph detection procedure which can identify an organized and biologically interpretable topological structure consisting of informative edges. The results in (d) are based on the ADSD method (see details in the Results section).

We propose a likelihood-based adaptive dense subgraph discovery (ADSD) model to extract informative connectomic subnetworks accurately. The new objective function is robust to edge-wise false positive and false negative noise by introducing a tuning parameter to balance the area density and degree density (Tsourakakis et al., 2013). We optimize the tuning parameter objectively by maximizing the widely used likelihood function in statistical network/graph model (e.g. stochastic block model, Holland et al., 1983, Zhao et al., 2012, Zhang et al., 2017). We develop efficient algorithms to implement the joint objective function of the ADSD. We further derive theoretical results which guarantee the approximation properties for any fixed graph and consistency for large graphs based on the proposed algorithm. We can perform permutation tests to yield statistical significance of extracted subgraphs (Ge et al., 2012, and Zalesky et al., 2010). We perform extensive simulation studies to validate the proposed model and theoretical conclusions. The results demonstrate improved accuracy of informative subgraph detection in various settings. Our method is then applied to a resting state fRMI (rfRMI) brain connectomic study for schizophrenia research. The results of our real data analysis reveal for the first time systematic aberrant salience network centered connectivity patterns in schizophrenia patients using whole brain connectome network analysis. Although some of our findings coincide with previous studies using seed voxel-based method, our analysis is more comprehensive and less biased, because it does not require pre-selected seed sets or focuses on exclusively known networks.

## 2 Methods

### 2.1 Background: group-level inference for multivariate edges in a graph space

Let *G* = (*V, E*) be an undirected graph, where 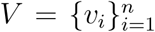 is a set of nodes representing brain areas and ROIs, and 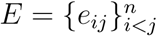 denotes the set of edges between pairs of nodes (i.e, connections between brain areas). *G′* = (*V′, E′*) is a subgraph of *G* if *V′ ⊂ V* and *E′ ⊂ E*. Then, *G*(*S*) = (*S, E*(*S*)) is a ‘nodes-induced’ subgraph if *S ⊂ V* and *E*(*S*) = {(*u, v*) ∈ *E*|*u, v ∈ S*} being edges in *E* with endpoints in *S*. We use 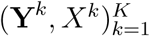 to represent the group-level multivariate edge data in a graph space *G* = (*V, E*), where *k* = 1, …, *K* is the subject index. 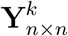 represents the brain connectome data in a binary/weighted adjacency matrix for subject *k*, and *X*^*k*^ is the corresponding vector of covariates (clinical and demographic variables). We assume that the location of nodes and edges are identical across subjects after spatial normalization. Thus, our goal is to perform statistical analysis and identify phenotype-related subnetworks with high sensitivity and well-controlled FDR (Lukemire et al., 2017, Kundu et al., 2018, Xia and Li, 2018, Vogelstein et al., 2012, Durante et al., 2018). Figure 1a demonstrates the procedure of group-level inference for brain connectome data.

Let 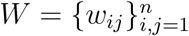 denote the edge-wise inference matrix based on graph *G*, where each off diagonal entry *w*_*ij*_ represents the edge-wise statistical inference results on edge *e*_*ij*_ (e.g., test statistics *t*_*ij*_ and *p* values *−* log(*p*_*ij*_)). For each edge *e*_*ij*_, we denote a corresponding latent indicator variable *δ*_*ij*_ such that *δ*_*ij*_ = 1 if edge *e*_*ij*_ is associated with the phenotype of interest and *δ*_*ij*_ = 0 otherwise. We consider the edge-wise inference results as our input data (Chen et al., 2020).

The goal of group-level brain network analysis is to identify a set of subnetworks 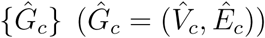 that are associated with a phenotype of interest, such that

1. 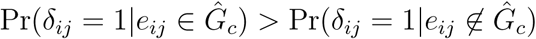 (dense subgraph);
2. The false discovery rate (FDR), 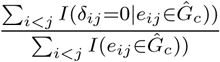 is low;
3. The sensitivity, 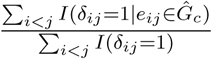 is high;
4. 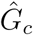 is a well defined community (e.g., a node-induced subgraph that 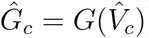.

In practice, the task above is challenging. For example, the mass univariate methods including both FDR and family-wise error rate (FWER) controlling models apply an universal threshold on all edges, and yield a set of unrelated ‘significant’ edges. Thus, they can neither address the trade-off between sensitivity and false positive findings by leveraging the information of network or yield findings with an organized and biologically interpretale network topological structure. The network based statistics (NBS) method allows edges borrow strengths from each other, yet it yields an unorganized subgraph (Zalesky et al., 2010, Fornito et al., 2012). Moreover, the signal subnetwork detected by NBS includes all nodes in *G almost surely*, i.e., *G*(*V*_*c*_) = *G*, when *n* is larger than a handful of nodes (Erdős and Rényi, 1960), and thus less interpretable.

We also notice that the proportion of true positive edges 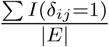 in *G* is often small in our motivated brain connectome data (e.g., around 5%), which may lead to the difficulty of applying the commonly used network models (Chen et al., 2018). Figure 1b shows the results of the application of stochastic block models (SBM) which miss the network topological structure. Therefore, it is highly desirable to extract a ‘dense’ subgraph which is a node-induced subgraph *G*(*S*_0_), such that the edge density *ρs* is much higher than the overall density *ρ*:

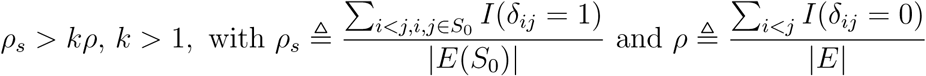

and *G*(*S*_0_) includes most edges. The detected informative subgraph can either directly become the subnetwork of interest or intermediate results for further refined network analysis (e.g., using SBM).

#### 2.1.1 Dense Subgraph Discovery

The conventional dense subgraph aims to detect a node-induced subgraph with maximized density. Two popular definitions of density function are also referred as average degree and edge ratio (Asahiro et al., 1996, Charikar, 2000, Lee et al., 2010):

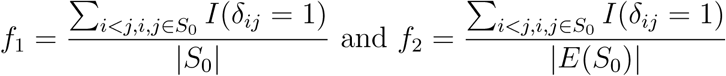

The edge ratio agrees with our goal for informative subgraph detection. However, the implementation of dense subgraph discovery is not trivial. The direct optimization of edge ratio *f*_2_ tends to detect a high-density subgraph with a tiny size (See Lemma 2 in supplementary for details). Meanwhile, it has been known the optimization of *f*_1_ can lead to the detection of an over-sized subgraph (Tsourakakis et al., 2013), which may cause a high false positive rate for statistical inference. Figure 1c shows the results of conventional dense graph discovery by optimizing *f*_1_. To address these challenges, we propose a likelihood based method for dense subgraph discovery.

### 2.2 Adaptive dense subgraph discovery

We consider *G* = (*V, E, W*) as our input data that stores edge-wise inference results in a weighted adjacency matrix *W*. Our goal is to extract a phenotype-related informative subgraph *G*(*S*) induced by nodes set *S* in the sense that *E*(*w*_*ij*_|*e*_*ij*_ ∈ *E*(*S*)) » *E*(*w*_*ij*_|*e*_*ij*_ ∉(*S*)) while maximally reducing false negative findings and improving the sensitivity.

To address the challenges in conventional dense subgraph discovery and improve the balance of the trade-off, we propose an adaptive density function:

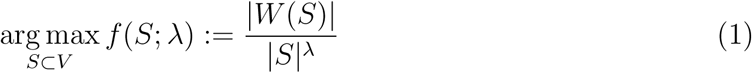

where *λ* ∈ [1, 2] is a tuning parameter, such that when *λ* = 1 and 2, *f* (*S*; *λ*) density function reduces to *f*_1_ and *f*_2_, respectively.

To better illustrate the impact of the tuning parameter *λ* on the FDR and sensitivity, we transform the objective function (1) to:

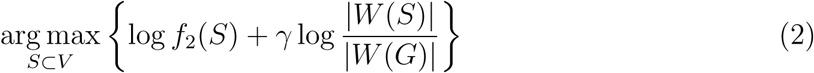

with *γ* ∈ [0, 1]. The optimal solution is approximated by *f* (*S*; *λ*) for large graphs with 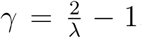. The first term in (2) is the true discovery rate (1 − FDR), while the second term is the sensitivity (power). In that, *λ* functions similarly to the tuning parameter in the shrinkage methods (e.g., LASSO) since *f*_1_ is related to the loss function and *f*_2_ implements the rule of parsimony. Increasing *λ* leads to a low FDR, while decreasing *λ* can improve the sensitivity. Therefore, our objective function is tailored for the four items of our overarching goal in the previous subsection.

In practice, both *G*(*S*) and *λ* need to be estimated, and *λ* is critical to balance the trade-off between FDR and sensitivity. We propose an iterative procedure to optimize the objective function (1) in subsection 2.2.1 and estimate *λ* in subsection 2.2.2. We name this new procedure adaptive dense subgraph discovery (ADSD).

#### 2.2.1 Optimization with a known *λ*

We implement the objective function (1) using a greedy algorithm. The greedy algorithm has been the most commonly used technique to implement objective functions for dense subgraph discovery (Asahiro et al., 1996, Charikar, 2000). Generally, a greedy algorithm removes a node with the minimum-degree at each iteration, and then selects the optimal dense subgraph from the process of node removal. The detailed procedure is described by Algorithm (3).

##### Algorithm 1 Optimizing objective function (2) with a given *λ*

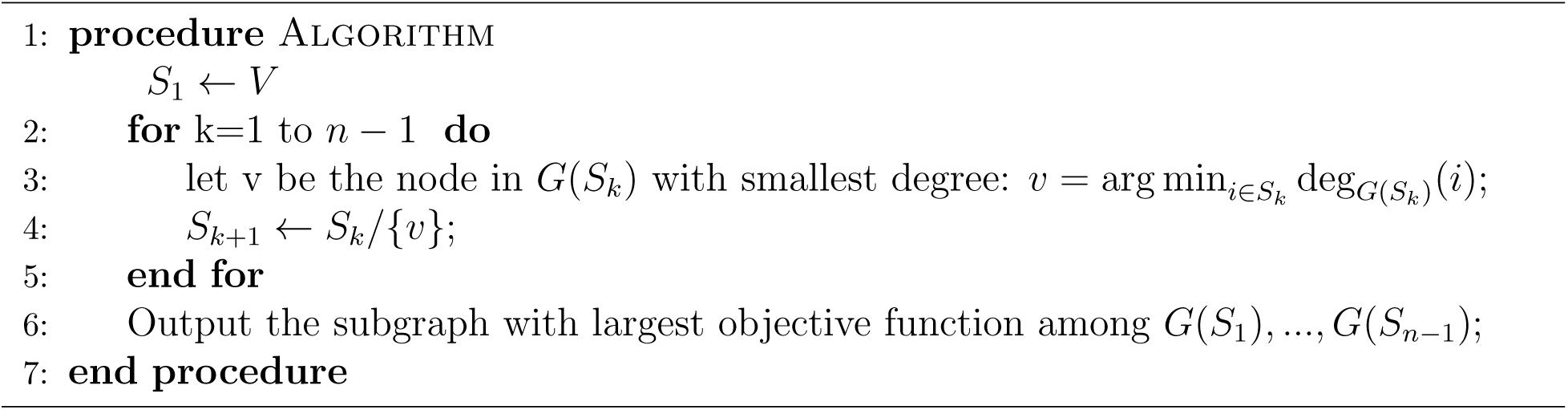

We denote the optimal dense subgraph based on our objective function (1) with a given *λ* by

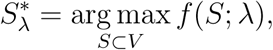

and the output of greedy algorithm as:

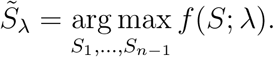

A major advantage of the greedy algorithm is the low computational complexity, which is critical for our application. Although our greedy algorithm may not provide the exact solution, Charikar (2000) proved the greedy algorithm has a 2-approximation as to *f*_1_(⋅), that is 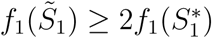, where 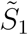 is the densest subgraph by greedy algorithm and 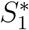 is the true maximizer for *f*_1_(⋅). In Section 3, we prove the theoretical approximation properties of 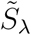 with regard to the maximization *f* (*S*^∗^; *λ*) for various values of *λ*.

#### 2.2.2 Likelihood-based method for *λ* estimation

Clearly, the performance of our greedy algorithm (3) relies on the unknown parameter *λ* (e.g. *λ* = 1 and 2 lead to the optimization *f*_1_(⋅) and *f*_2_(⋅) alone respectively). We propose a data-driven approach to automatically determine *λ* by maximum likelihood estimation. In statistical literature, the likelihood function of network/graph data has been well studied (Zhang et al., 2017). For example, a binary graph with *K* block can be defined by:

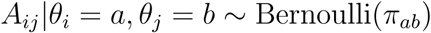

where *A*^*n×n*^ is a binary adjacency matrix, ***θ*** = (*θ*_1_, …, *θ*_*n*_) is a latent vector of node labels, and 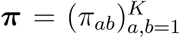 is a *K* × *K* symmetric probability generating matrix for generative for edges within and between blocks/communities.

We adopt the likelihood function of SBM because the dense subgraph structure in our ADSD model can be considered as a special case of the block diagonal structure in SBM. Specifically, in our model the graph *G* = (*V, E*) includes an underlying true informative subgraph *G*(*S*0) and all other nodes are singletons. The number of communities of SBM is *K* = *n* − *n*_*s*_ + 1, where *n* = *|V|* and *n*_*s*_ = |*S*_0_|. We further assume the planted partition model that the parameters of Bernoulli distributions for edges between blocks are identical in SBM.

To construct the likelihood function for ADSD, we first binarize the input data matrix *W* using a threshold *r* and let *A*_*ij*_ = {*W* (*r*)}_*ij*_ = *I*(*W*_*ij*_ > *r*). We denote ***θ***(*S*) as a vector of node labels concerning the node set *S* for a dense subgraph *G*(*S*), where an element *θi*(*S*) = 1 if *i* ∈ *S* and *θi*(*S*) = 0 for *i* ∈ *V/S*. Then, the membership of edges regarding the nodes-induced subgraph *G*(*S*) can be defined consequently as *θ*_*ij*_(*S*) = *θ*_*i*_(*S*)*θ*_*j*_(*S*).

We let all edges in *A* follow a Bernoulli distribution with parameters *π*_*ij*_ that

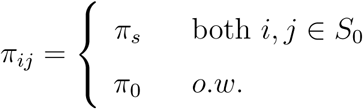

Using the mixture model representation, *π*_*ij*_ = *θ*_*ij*_(*S*_0_)*πs* + (1 *− θ*_*ij*_(*S*_0_))*π*0. The likelihood function based on *A* = *W* (*r*) is:

When the membership of informative subgraph is given, the MLE of the edge probabilities can be obtained by:

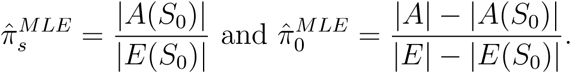

In practice, *S*_0_ is unknown and can be estimated by 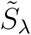 from the Algorithm (3) with a given *λ*. The likelihood function based on 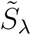 is in the form:

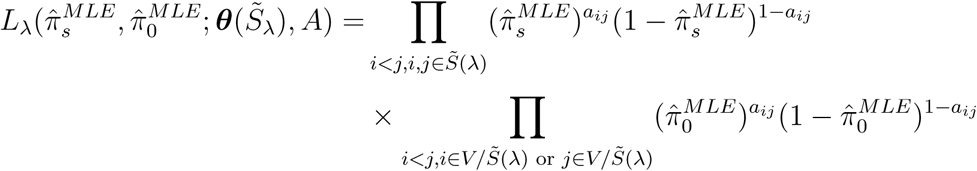

where 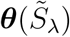 is the node label vector associated with 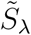.

Therefore, *λ* can be estimated by two steps. First, for any *λ* ∈ [1, 2], we can extract a dense subgraph 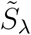 by the greedy algorithm (3). Next, 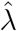 is determined by the combination of *λ* and 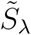 that maximizes the likelihood function:

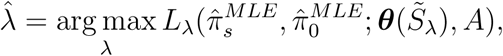

The final result of dense subgraph discovery based on the MLE determined 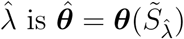.

We further consider the threshold *r* in *A*_*ij*_ = {*W* (*r*)}_*ij*_ = *I*(*W*_*ij*_ > *r*) as a random variable following a distribution *g*(*r*) rather than a fixed value in order to avoid an arbitrary selection. We integrate the likelihood with respect to *r* based on the prior distribution *g*(*r*), and thus our optimization is invariant to the selection of *r*. *g*(*r*) can be a discrete distribution with a support {*r*_1_, …., *r*_*m*_} and corresponding probability {*g*(*r*_1_), …, *g*(*r*_*m*_)}. In practice, the performance of our algorithm is robust to the prior distribution, given the reasonable support of *r* is used. By integrating *r* out, the likelihood function becomes:

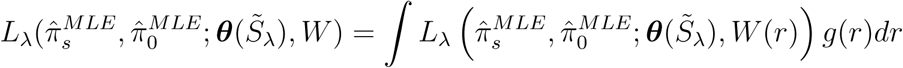

The general algorithm for ADSD is described in the Algorithm (2). Since Algorithm (3) is nested within the overall Algorithm (2), the low computational cost of Algorithm (3) is critical for the overall computational efficiency of ADSD. The complexity of the ADSD algorithm is *O*(*M n*^2^) where *M* is a sufficient searching range of *λ*. The resulting subgraph 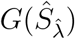 from our ADSD model can be further investigated for more delicate latent topological structures and statistically tested by permutation tests with family-wise error rate control (Zalesky et al., 2010, Chen et al., 2015) and we include the details in the supplementary materials.

##### Algorithm 2 The complete ADSD algorithm

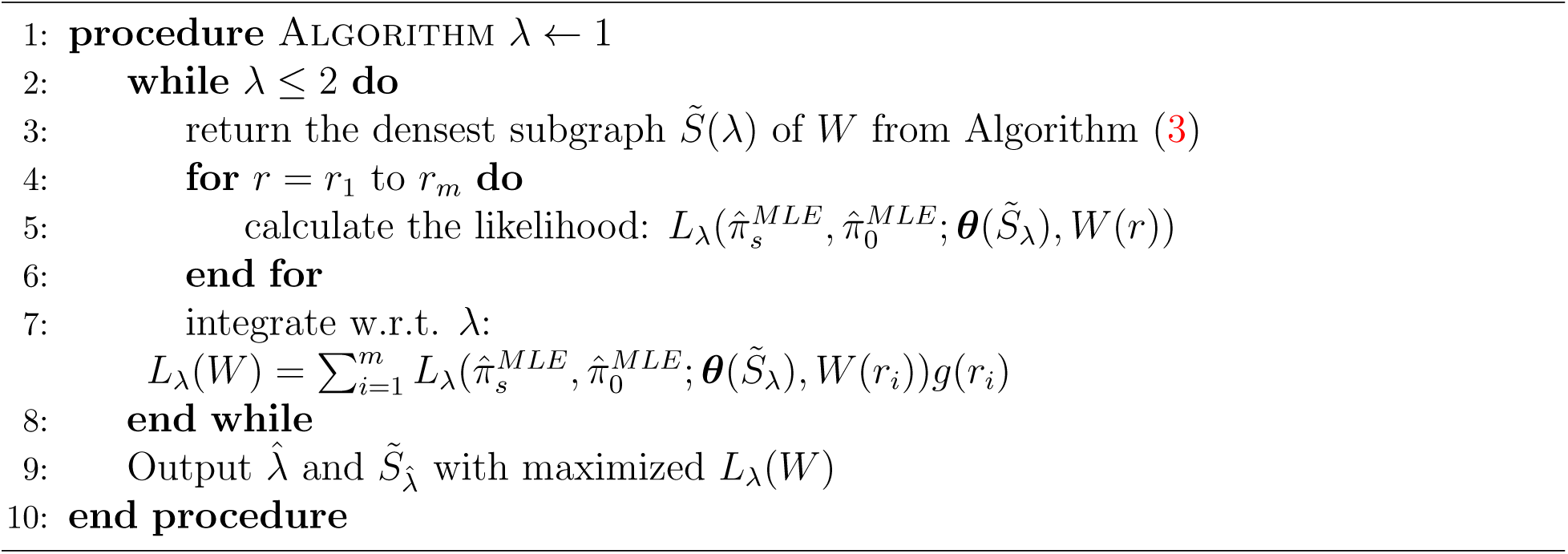

## 3. Theoretical results

The theoretical work for conventional dense graph discovery has been well-established (Lee et al., 2010). For example, Charikar (2000) showed that the commonly used greedy algorithm proposed by Asahiro et al. (1996) has a 2-approximation bound. In this article, we aim to extend the theoretical results for our new ADSD algorithms in 2 which generalizes the traditional objective function by introducing the parameter *λ*. Specifically, we discus the approximation bounds for ADSD with a full range of *λ* values in the following theorem 1.

### Theorem 1 (Exact property of Algorithm 3)

*For a given graph G* = (*V, E*), *with 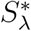 and 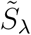 defined in section 2, the Algorithm (3) has a ρ*(*λ, n*)*-approximation, especially* 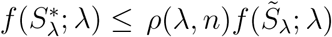 *with*

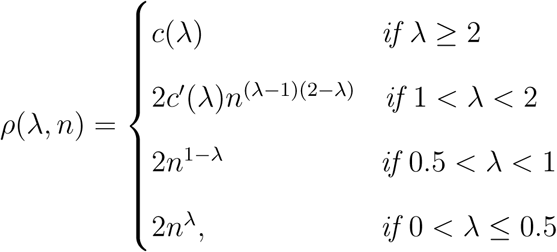

where *c*(*λ*) = 2^λ−1^ and *cI*(*λ*) = 1 *∨* 2^1−λ^.

The theorem 1 provides the performance of Algorithm (3) by guaranteeing the closeness of objective function in 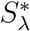 and 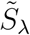. However, an optimal optimization may not result from a perfect recovery of informative subgraph for randomness (i.e. 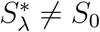 for all *λ*). Hence, we further prove the asymptotic consistency 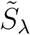 from Algorithm (3) in following theorem 2. When the observed graph is generated from some underlying model with true informative subgraph *S*_0_, there exist an *λ* such that 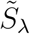 tends to *S*_0_ with probability 1 asymptotically.

### Theorem 2 (Asymptotic property of Algorithm 3)

*Assume the graph G = (V, E) including an informative subgraph G*(*S*_0_) = (*S*_0_, *E*(*S*_0_)) *is generated from the special SBM we defined in section 2.2.2, such that the edges are drawn from independent Bernoulli distributions with parameter *π*_*ij*_ = *π*_*ij*_(*S*_0_) = *θ*_*ij*_(*S*_0_)*πs* + (1 − *θ*_*ij*_(*S*_0_))*π*_0_, where *θ*_*ij*_(*S*) = *θ*_*i*_(*S*)*θ*_*j*_(*S*), *θ*_*i*_(*S*) = *I*(*i ∈ S*) *and πs > π*0*. Let |S*0| = *O*(*|V|*1/2+ϵ*) *as n → ∞ for any ϵ* > 0.

*Then, there exist some λ such that we will get exact recovery with probability 1 in Algorithm (3), i.e. as n → ∞*,

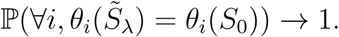

Theorem 2 provides the existence of parameter *λ* for a consistent estimator as the size of graph goes to infinity. We use the following Theorem 3 to demonstrate the performance of Algorithm (2) by illustrating the selected *λ* based on our likelihood-based criterion will lead to an estimator with negligible proportion of incorrect assignment for nodes.

### Theorem 3.

*Assume the graph G = (V, E) includes an informative subgraph G* (*S*_0_) = (*S*_0_, *E*(*S*_0_)), *such that the edges generate from independent Bernoulli distributions with parameter π*_*ij*_ = *π*_*ij*_(*S*_0_) = *θ*_*ij*_(*S*_0_)*πs* + (1 − *θ*_*ij*_(*S*_0_))*π*_0_, where *θ*_*ij*_(*S*) = *θ*_*i*_(*S*)*θ*_*j*_(*S*), *θ*_*i*_(*S*) = *I*(*i ∈ S*) *and πs > π*0. *Let |S*0| = *O*(*|V|*1/2+*c*) *as n → ∞ for any ϵ >* 0.

*Then, as n → ∞, the adaptive greedy algorithm with likelihood-based criterion results in an estimate* 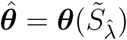 *with*:

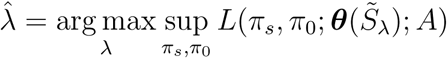

*has incorrect assignment with probability converging to zero, i.e*.

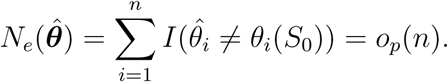

The detailed derivations and proofs for the above three theorems are provided in the supplementary materials.

## 4 Results

We apply the proposed ADSD method to the neuroimaging data collected from patients with schizophrenia and healthy controls. This data set includes 104 patients with schizophrenia (SZ) (age 36.88 *±* 14.17, 62 males and 41 females, 1 other) and 124 healthy controls (HC) (age 33.75 *±* 14.22, 61 males and 63 females). There are no systematic differences in age (test statistic 1.64, *p* value 0.10) or gender (test statistic 1.67, *p* value 0.10) between the two groups. The imaging acquisition and preprocessing details are described in Adhikari et al. (2019). A brain connectivity-based atlas is used to denote 246 regions of interest (ROIs) as nodes in a brain connectome graph (Fan et al., 2016). The functional connection (edge) between a pair of nodes for each subject is calculated by the covariation between averaged time series from the two corresponding brain ROIs. The Fisher’s Z transformed Pearson correlation coefficient then is applied for each edge. We perform non-parametric group level testing on each edge, although alternative inference methods can be used as well.

We focus on the input matrix *W* reflecting the importance levels (− log(*p*_*ij*_)) on all edges, as demonstrated in Figure 2a. We first apply the greedy algorithm (e.g., Charikar’s method that is equivalent to the proposed greedy algorithm with an ad-hoc *λ* = 1) for dense subgraph extraction. The results in Figure 2b seem to be an over-inflated subnetwork without clear biological interpretation and a large set of false positive edges. We also applied other popular subgraph detection methods, for example, breadth first search in network-based statistics, stochastic block model, and various community detection methods (Zalesky et al., 2010, Amini et al., 2013, Newman and Girvan, 2004). However, these algorithms either detect a subgraph including all brain regions or yield no findings. In contrast, by implementing our ADSD method (2), we obtain a subnetwork 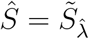 with 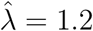. We note that the detected subgraph is robust to the prior distribution of *G*(*r*) as long as a reasonable support is used.

**Figure 2:**
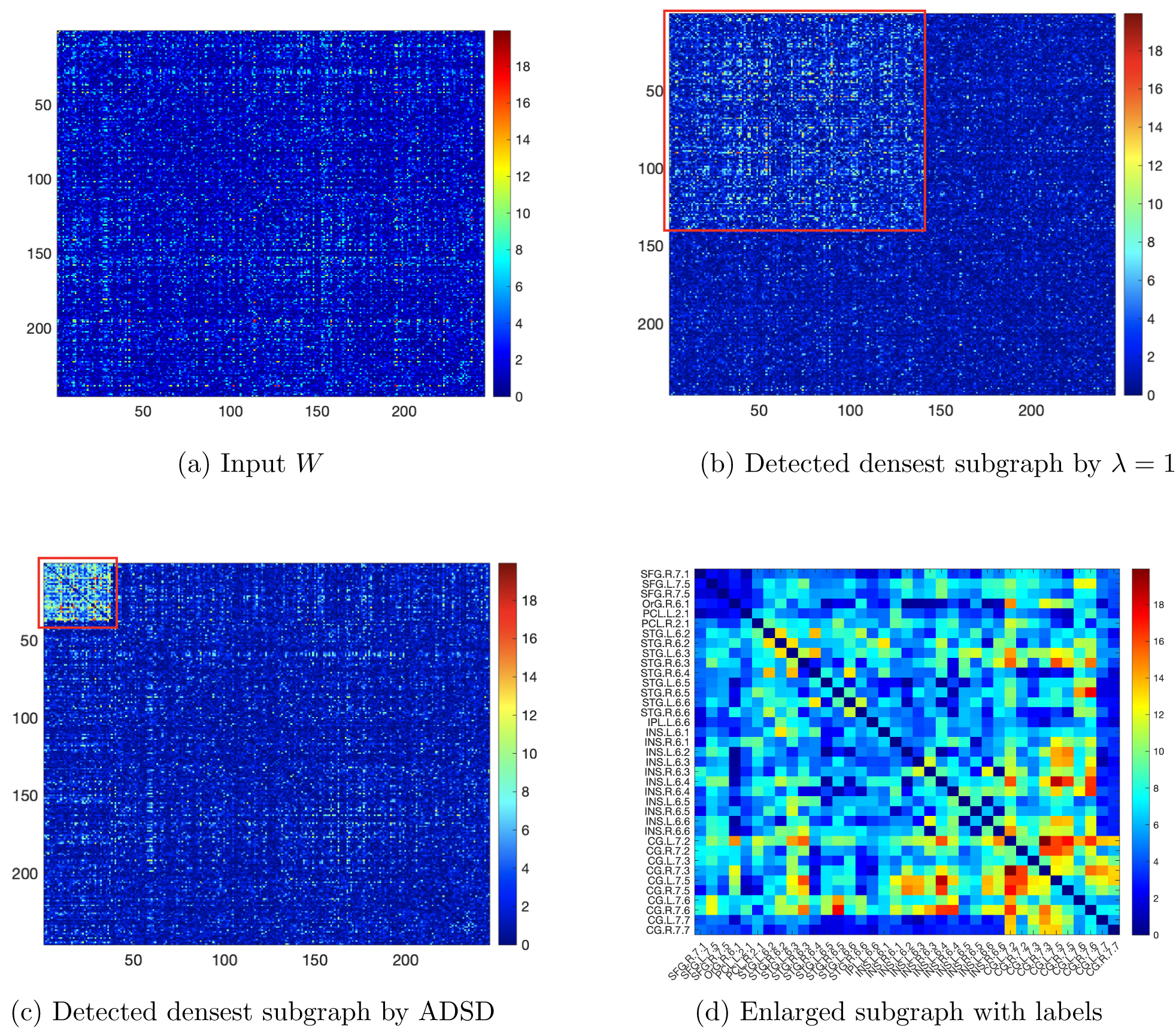
Results of data example: (a) is the input matrix *W*; (b) shows the results of existing dense graph discovery; (c) demonstrates the results by applying ADSD; (d) illustrates refined topological structure based on results of ADSD.

The computation is efficient, and it takes 2.21 seconds to implement the ADSD algorithm on a Mac with CPU Core i5 and memory 8GB. We further calculate the p-value of the network based on permutation test (Zalesky et al., 2010, Chen et al., 2015, Chen et al., 2020). The p-value for the network is significant *p* < 0.001 with family wise error rate adjustment.

The results show a subnetwork with reduced functional connectivity in patients with schizophrenia compared to healthy controls (see in Figures 3), which is consistent with the current knowledge that schizophrenia is possibly a degenerative disorder and associated with hypoconnectivity (Li et al., 2019). This subnetwork is centered around the well-known salience network (SN) which is primarily composed of bilateral insular gyri (INS) and anterior cingulate cortices (ACC). The salience network contributes to complex and integrative brain functions including emotions, cognition, and self-awareness (Uddin, 2015). Numerous previous studies have reported that decreased functional connectivity in the salience network is related to several core symptoms of schizophrenia using seed voxel methods (Palaniyappan et al., 2012). Our findings are well aligned with these established results. In addition to SN, our subnetwork extracted by ADSD involves several other brain regions including bilateral superior temporal gyri (STG), superior frontal gyri (SFG), precentral gryi (PCL), inferior parietal lobe left (IPL), and orbitofrontal cortex right (OrG). These regions have been identified to associate with auditory perceptual abnormalities (STG, IPL), voluntary movement (PCL), and sensory and cognition (SFG, OrG) (Fornito et al. (2012)). Jointly, our detected subnetwork reveals a comprehensive and systematic brain connectivity aberrance in patients with schizophrenia, which is related to the impaired capability to integrate and comprehend information (e.g., multiple external stimuli) and to respond appropriately. The detected schizophrenia-related brain connectome subnetwork is biologically plausible. It provides evidence to combine prior isolated findings, and thus enhances our understanding of the complex brain connectomic patterns and clinical symptoms.

**Figure 3:**
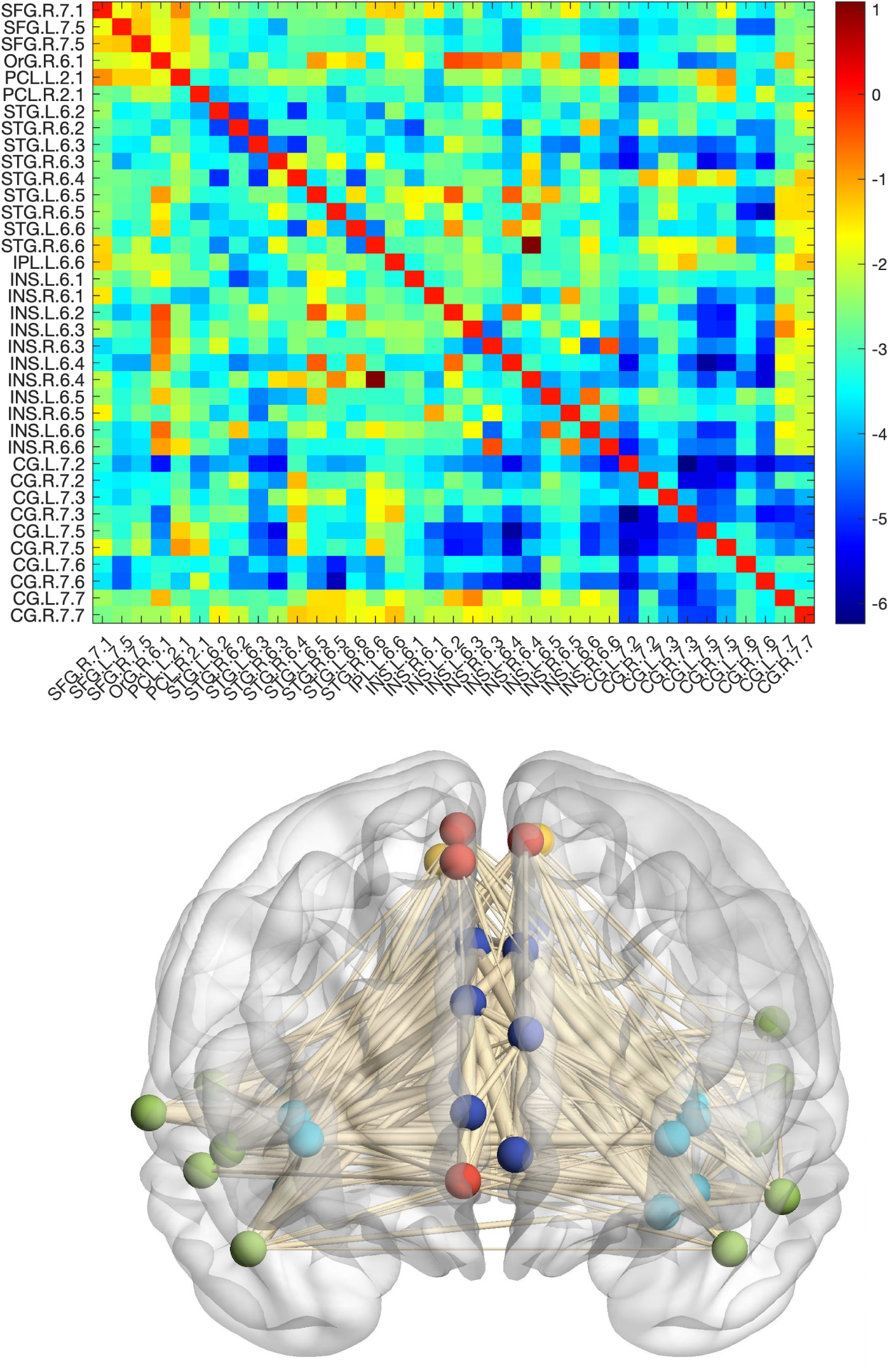
(a) illustrates the enlarged and labeled informative subgraph in t-statistics detected by ADSD which indicates decreased functional connectivity of SZ. (b) is a 3D demonstration of the subgraph: red nodes represent superior frontal gyrus (SFG) + orbitofrontal cortex right (OrG); yellow nodes are precentral gryi (PCL); green nodes are superior temporal gyrus (STG)+inferior parietal lobe left (IPL); blue nodes represent insular gyrus (INS); navy nodes represents cingulate cortex (CG).

Thus, our novel analytic approach revealed a neural sub-network that has been previously shown to both differentiate healthy controls and patients with schizophrenia and has been critically linked to core symptoms of the disorder. Since our results do not depend on the arbitrary selection of seed voxels and pre-specified networks of interest, our results are subject to less selection bias and thus more reliable and comprehensive.

## 5. Simulation

In the simulation study, we generate multiple brain connectivity data sets under several settings. We consider a graph *G* with *|V|* = 100, where an informative subgraph in a community structure with two possible sizes *|S*_0_| = 15 and 30. We generate 60 connectivity matrices for 30 controls and 30 cases. We assume that most edges in the informative subgraph are differentially expressed between cases and healthy controls. We let the connectivity weights of edges inside the informative subgraph follow a normal distribution with mean *µ*_1_ and variance *σ*^2^, while all other edges have normal *µ*_0_ and *σ*^2^ for the case group. In the control group, we let all edges follow a normal distribution of *µ*_0_ and *σ*^2^. Specifically,

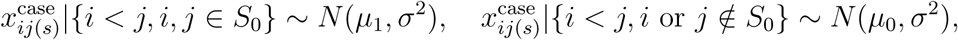

and

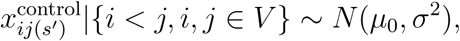

where 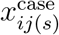 represents the edge linking node *i* and *j* for the *s*th subject in case group, and 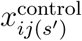 control defines the edge weight for the *sI*th subject in control group.

We apply various standard effect sizes (i.e., signal-to-noise ratios - SNRs) by setting *σ* = 1, and *µ*_0_ = 0, *µ*_1_ = 0.4, 0.6, 0.8. We further consider a more realistic scenario by letting the proportion *q*_1_ of edges inside informative-subgraph be non-differentially expressed (i.e. *N* (*µ*_0_, *σ*2) for both cases and controls). Similarly, we set a *q*_2_ proportion of edges outside informative-subgraph are differentially expressed (i.e. *N* (*µ*_1_, *σ*^2^) for cases and *N* (*µ*_0_, *σ*^2^) for controls). (*q*_1_, *q*_2_) represent the practical non-perfect distribution of informative edges in the overall graph. In the simulation data, two sets of parameters (*q*_1_, *q*_2_) = (0.8, 0.1) and (0.9, 0.05) are used.

We compare the ADSD method with the two most popular dense subgraph discovery methods including Greedy algorithm with *λ* = 1 and Goldberg’s algorithm. The results are evaluated by node-assignment accuracy in terms of true positive rate (TP) and true negative rate (TN) defined as follows:

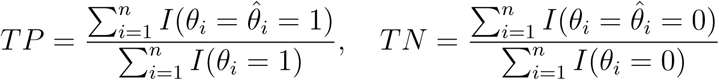

The mean and standard errors of TP and TN for three methods across 30 replicates for all settings are displayed in the following Tables 1 and 2.

**Table 1:**
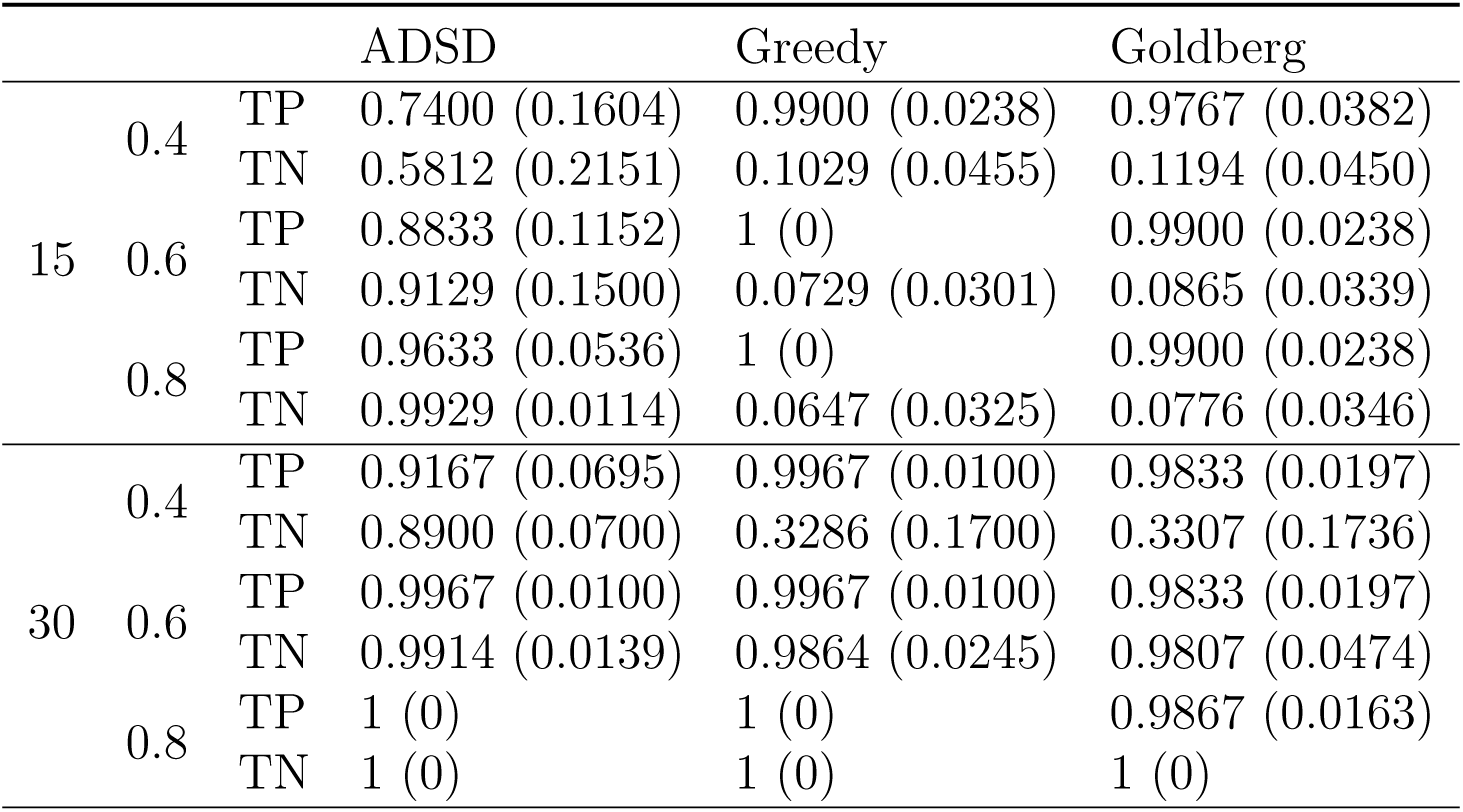
The node-assignment accuracy of three methods under varied SNRs and subgraph size with (*q*_1_, *q*_0_) = (0.8, 0.1)

**Table 2:**
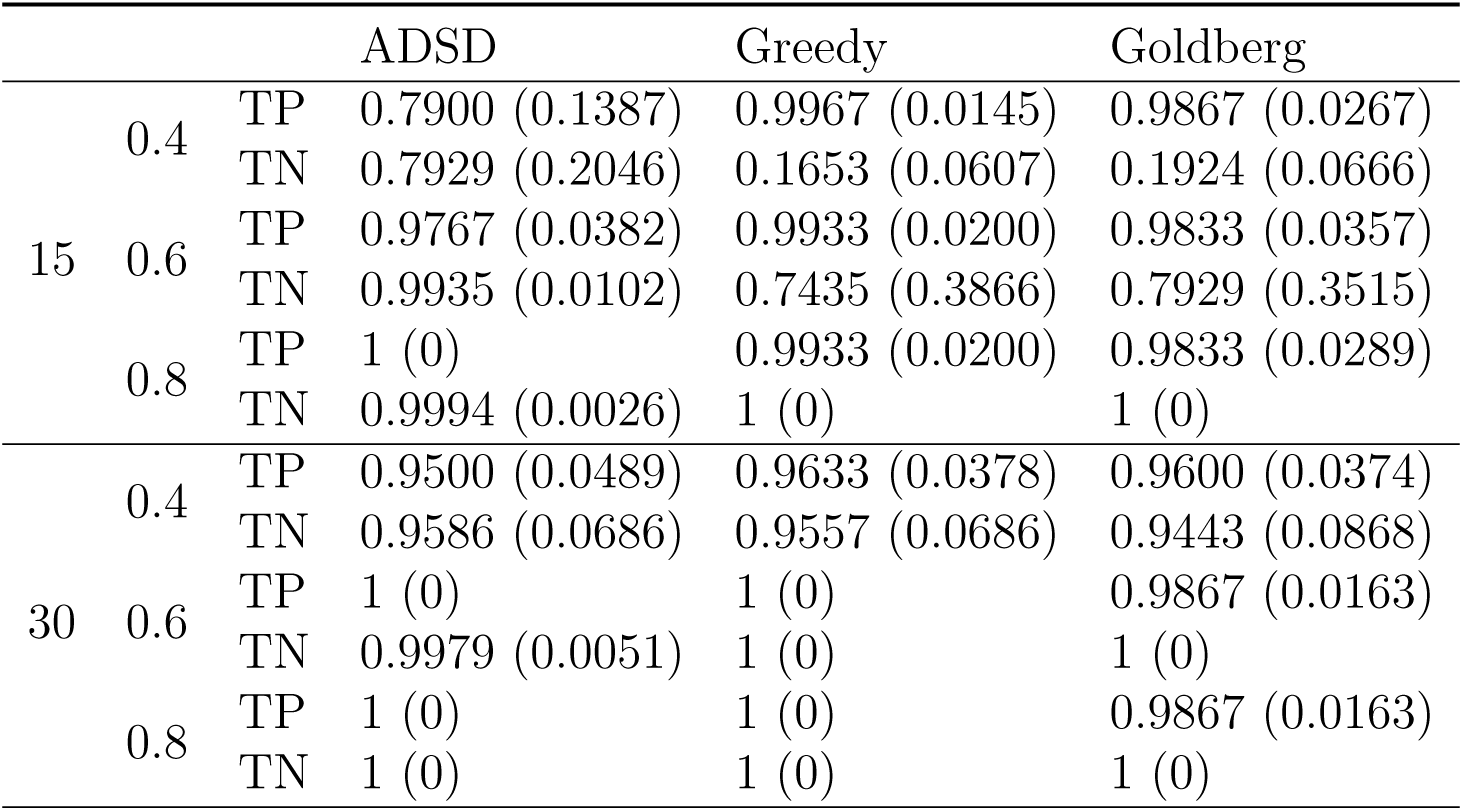
The node-assignment accuracy of three methods under varied SNRs and subgraph size with (*q*_1_, *q*_0_) = (0.9, 0.05)

Table 1 demonstrates results with (*q*_1_, *q*_0_) = (0.8, 0.1). When the subnetwork size is 15, the Charikar’s greedy algorithm and Goldberg’s algorithm have extremely high values of true positive rate and low true negative rate for all small and large effect sizes/SNR. This indicates that the conventional algorithms tend to detect over-sized dense subgraphs consisting of a high proportion of false positive edges when the subgraph is small and SNR is medium-large. This phenomenon is common in brain connectome data analysis in practice: the size of disease-related subnetwork is relatively small (the proportion of non-null distribution is small) with medium edge-wise effect sizes which can be disturbed by false positive and positive signals. If a universal threshold is used for all edges (e.g., BH-FDR and FWER cutoffs), then the most true positive edges will be missed along with the topological structure due to the control of false positive findings. In contrast, the proposed ADSD approach is robust to the noise and can accurately extract the informative subgraph. With increasing strengths of signal (i.e., larger informative subnetwork size and SNR), the traditional dense graph discovery algorithms are comparable to ADSD and they all outperform edge-wise inference (e.g., FDR and FWER). Table 2 shows the results with higher accuracy of subnetworks (*q*_1_, *q*_0_) = (0.9, 0.05). The ADSD can more accurately recover the dense subgraph when the SNR is small-medium and robust to false positive and false negative edges. Similarly, all three methods show comparably perfect performance when the subnetwork size and/or SNR are large.

In summary, the simulation results clearly show that likelihood-based ADSD approach is more robust to both false positive and false negative noise and can better capture smaller subnetworks with a high sensitivity and a low false positive rate. These properties are critical for the brain connectome analysis in practice because the real data sets are often mixed with substantial noise and include a small proportion of signal edges.

## 6 Discussions

In this article, we compare brain connectome matrices between diagnostic groups (e.g. schizophrenia and healthy subjects) to understand connectivity patterns altered by psychiatric illness. As in our motivation data example, however, disease-related subnetworks can be overwhelmed by substantial noise in the connectome data and thus difficult to extract. The noise heavily influences statistical inference by introducing enormous edge-wise false positive and negative errors that are constrained in a weighted adjacency matrix, and thus impose difficulty in understanding the network topology of disease-related brain circuits and in yielding valid statistical inference.

To overcome these challenges, we develop a novel ADSD method to reliably and robustly identify signal subgraphs (related to the phenotypes of interest) from the whole brain connectome network. The overall brain connectome inference network is often over-sized with a small proportion of signal edges which are not compatible with existing statistical network models. Therefore, it is desirable to detect a dense subnetwork maintaining most signal edges in a clique with a much smaller number of nodes (nodes induced subnetwork) and discarding a large proportion of false positive edges from the overall network. Dense graph discovery has been a popular research topic in network analysis for a couple of decades. Dense graph discovery methods are distinct from existing statistical methods for network analysis (e.g. various versions of community detection) because they focus on a network with a far fewer number of connections than a highly connected network consisting of communities. The dense graph discovery method is well suited for our application because the number of edges from the non-null distribution is relatively small (Efron, 2012). A key limitation of the current dense graph discovery methods is sensitive to noise. Due to the substantial noise in brain connectome data, the existing dense graph discovery methods tend to extract over-sized dense subgraphs which can lead to a high FDR, potentially incorrect biological findings, and low replicability. The proposed ADSD method integrates the concept of shrinakge into dense graph discovery by introducing a balance parameter to include the most informative edges into the subgraph (high sensitivity) while maintaining a low FDR. The balance parameter can be estimated based on the likelihood function which is commonly used in network statistics. We develop efficient algorithms to implement the objective function that is compatible with computationally intensive inference methods (e.g., permutation tests and bootstraps). In the current research, we apply permutation test based statistical inference on the dense subgraph. Both the simulation and data example results show that the proposed method is robust to the false negative and positive edges and can accurately detect the target dense subgraph with high sensitivity and low false positive rates. Therefore, our goal of brain connectome analysis can be well met by applying ADSD.

Our work makes several contributions to the field: first, the ADSD objective function and algorithms provide new dense subgraph detection tools for noisy, weighted, large, and less dense graphs, which may have wide applications in data mining and knowledge discovery. Secondly, we derive theoretical results to provide the bounds for the approximation of ADSD algorithms in a full range of the balance parameter. The asymptotic property of subgraph detection and balance parameter estimation are also developed. Last, the biological findings are novel, integrative, and clinically meaningful. Although part of these findings has been found in previous studies, only edge-wise results (i.e. links between regions to a fixed seed) are reported without fully investigating the interactive nature of network-level inference.

In this article, the hypo-connections in the salience network centered subnetwork groups in patients with schizophrenia are detected for the first time by whole brain connectome network analysis with explicit network topology. The reported network reveals the novel links between aberrant functional connectivity networks and impaired capability to integrate information from multiple sources (cognition deficits) in patients with schizophrenia, which may assist to further understand the underlying biological mechanism for multiple schizophrenic disorder symptoms.

In summary, we develop a likelihood-based adaptive dense graph detection method to extract the dense subgraph from a large and noisy network (weighted and/or binary). Our ADSD method outperforms existing dense subgraph discovery methods when the overall graph includes a small proportion of edges with high importance levels, and thus is well-suited for group-level brain connectome analysis. ADSD can also serve as a screening step for group level network analysis to effectively extract a dense subnetwork from a large overall network for further analysis. In addition, ADSD can be applied to other biological network data (e.g. interactive networks of genomics and proteomics data) and yield findings revealing latent and complex co-expression subnetworks. Therefore, ADSD can become a new useful tool for statistical analysis of large and less dense networks.

## 7 Appendix A

### 7.1 Permutation Tests

We could evaluate the significance of detected subgraph by permutation testing (Zalesky et al., 2010). For an undirected graph *G* = (*V, E*), the edge permutation permutes the order of edges. The edge-permuted graph according to a permutation *π* is given by *G*_*π*_ = (*V, π*(*E*)), with {*e*_*ij*_ ∈ *E}* = {*e*_*π*(*i*)*,π*(*j*)_ ∈ *E*_*π*_}. For the randomness of permutation, we have *{e*_*ij*_ ∈ *E*} ⊥ {*e*_*ij*_ ∈ *E*_*π*_}. Therefore, the organized pattern of *G* does not exist for *G*_*π*_, and the edge-permuted graph *G*_*π*_ becomes a random graph with probability being the proportion of connected edges in *G* (i.e. 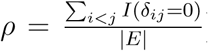). Hence, the significance of the topological *|E|* pattern of *G* can be evaluated among random graphs with identical proportion of connected edges *G*_*π*_.

In a weighted graph *G* = (*V, E, W*) with *|V|* = *n*, let *φ*(⋅) be the half-vectorization of a symmetric matrix. Denote *π* as a permutation of 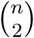 elements, and *P*_*π*_ is the corresponding permutation matrix. Then, the weight matrix of the edge-permuted graph *G*_*π*_ = (*V, E*_*π*_, *W*_*π*_) is given by *W*_*π*_ = *φ*^−1^(*P*_*π*_*φ*(*W*)).

A true informative subgraph includes edges with higher average level of importance (i.e. p-values) which can not be extract from a random graph. Hence, we propose a network level statistic such that it prefers subgraphs with more informative edges (larger subgraphs that can not be generated by random) and averagely higher level of significant edges (smaller subgraphs by excluding less non-significant edges). For a graph *G* with informative subgraph *G*(*S*), let

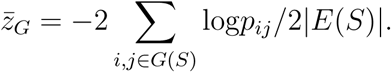

We propose the test statistic based on Fisher’s combination test and Chernoff bound of *χ*^2^ the cumulative distribution function:

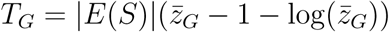

The whole procedure of the permutation testing is as follows:

#### Algorithm 3 Permutation testing of extracted subgraph

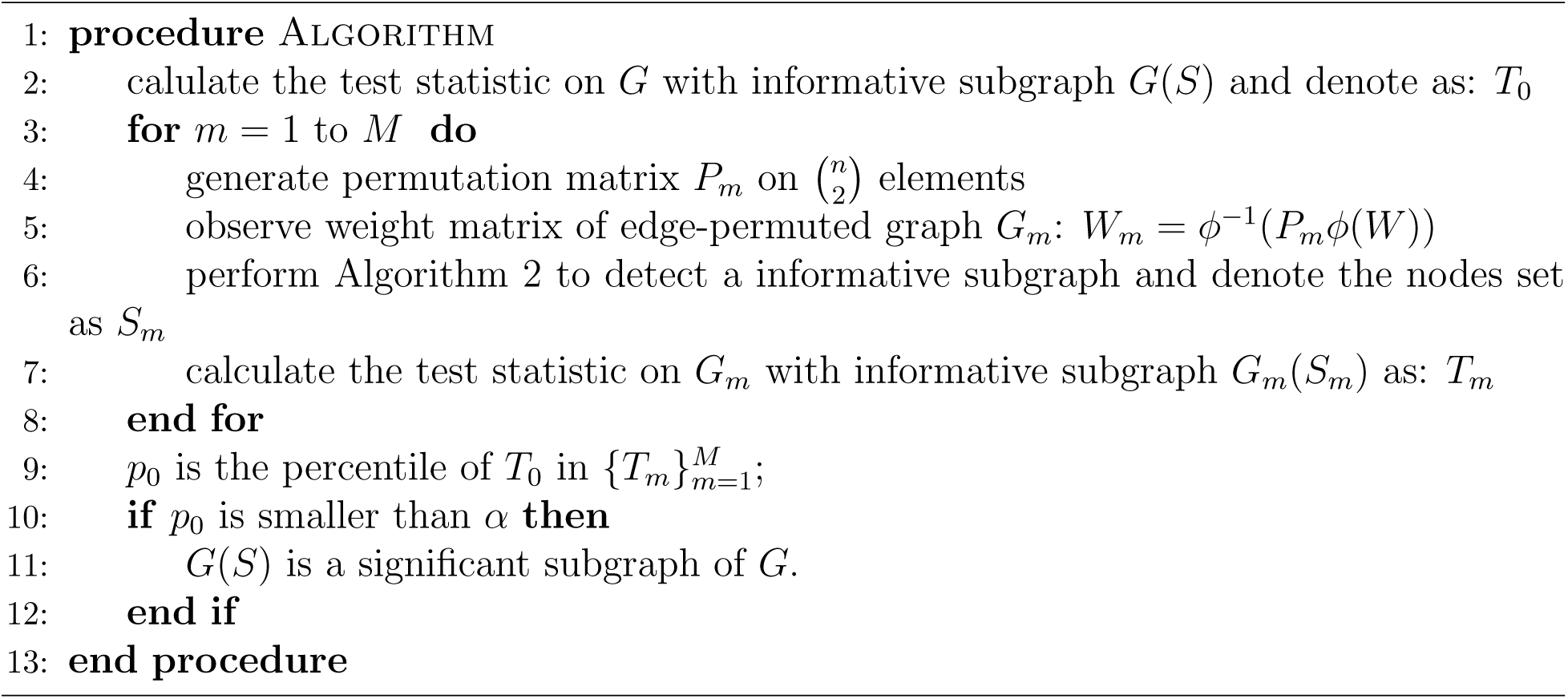

#### 7.2. Related work in densest subgraph

Due to noise, dense graph discovery algorithms tend to either identify over-sized subgraphs that may include a large proportion of false positive edges with low importance levels (high false positive discovery rate) or detect over-conservative small-sized subgraphs that may not sufficiently cover edges with high importance levels (low sensitivity, Tsourakakis et al., 2013). The two following lemmas are constructed to demonstrate the cases where conventional dense discovery methods may lead to detecting trivial-sized subgraphs. The generative mechanism is considered to be the same as stated previously.

##### Lemma 1

(Densest subgraph regarding average degree *f*_1_). *Assume that a graph G* = (*V, E*) *with an induced dense subgraph G*(*S*_0_) = (*S*_0_, *E*(*S*_0_)) *is generated by the same joint distribution as Theorem 3 with n*^1/2+*ϵ*^ ≲ *n*_*s*_ ≤ *cn*. *Let π_s_ > π*_0_ *> ϵ*_1_ *and cπ*_0_ *− c*_2_(*π_s_ − π*_0_) *>* 2*c*_1_*n^−ε^ be some positive constant ε*_1_ *and c*_1_*. Then, denoting the densest subgraph as 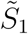, we have*

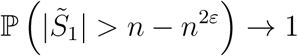

##### Lemma 2 (Densest subgraph regarding edge ratio *f*_2_)

*Assume that a graph G* = (*V, E*) *with an induced dense subgraph G*(*S*_0_) = (*S*_0_, *E*(*S*_0_)) *is generated by the same joint distribution as Theorem 3 with n*^1/2+*ϵ*^ ≲ *n_s_ ≤ c_n_. Let πs > π*_0_ *> ϵ*_1_. Then, for the densest subgraph 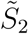, *we have*

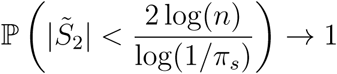

*almost surely as n* → ∞.

Note for lemma 2, the upper bound is relatively small compared with the size of graph. For example, for *n* = 1000 and *π*_*s*_ = 0.5, 2 log *n/* log(1*/πs*) *≈* 19.93, and for *n* = 10000 and *π*_*s*_ = 0.7, 2 log *n/* log(1*/π*_*s*_) *≈* 51.65. The proofs for these two lemmas are included in the appendix. Therefore, the direct application of conventional dense subgraph discovery algorithms to our network data may miss the underlying important findings of disease-related subnetworks.

#### 7.3 Proofs

Before we prove the Theorem 1, we first establish the following Lemma 3 by applying the idea of proof in Charikar (2000) with density function *f*_1_(*S*) directly.

##### Lemma 3.

*The greedy algorithm will give a* 2*n*^|*λ*−1|^ *approximation, i.e. f* (*S∗*) *≤* 2*n*^|*λ*−1|^*v*.

*Proof*. For an undirected graph, we assign the edge *e*_*ij*_ to either node *i* or *j*. Denote the number of edges assigned to node *i* as *d*(*i*). Then,

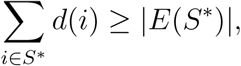

since all edges in *E*(*S∗*) will be assigned to node *i* or *j* which will be included in Σ_*i∈S∗*_ *d*(*i*).

Consider a specific way to assign edges. Let the node gets assigned when it is removed in the greedy algorithm. In other words, *d*(*i*) equals the degree of node *i* in the iteration that it is removed in greedy algorithm. Assume the subgraph at the iteration that *i* is removed is *G*(*S′*) such that at this iteration the nodes set changes from *S′ → S′/{i}*. Therefore, since the node *i* is deleted for the smallest degree, we will have the degree of *i* is smaller than the *average degree in this iteration*:

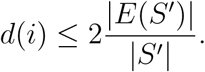

**Case 1:** For *λ <* 1,

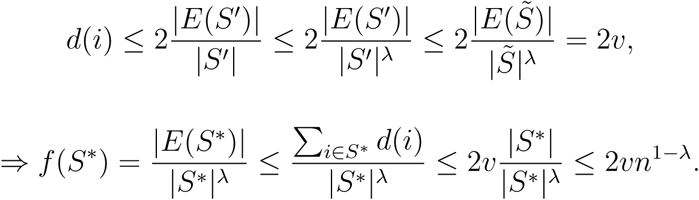

**Case 2:** For *λ >* 1,

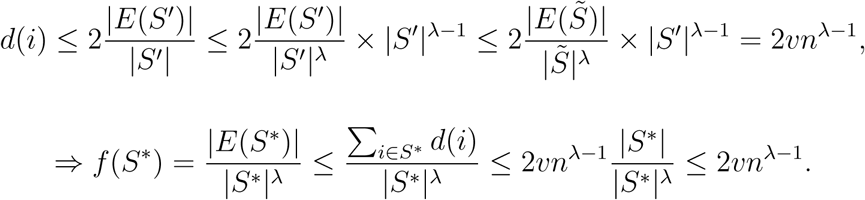

##### 7.3.1 Proof of theorem 1

**Case 1:** *λ ≥* 2. The subgraph with only two nodes has *f* (*S*; *λ*) = 1/2^*λ*^, otherwise all elements in this 2 *×* 2 matrix should be zero and inductively the raw graph have no edges (all elements in the corresponding matrix should be zero). Thus, 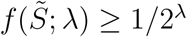.

On the other hand, when *λ* ≥ 2,

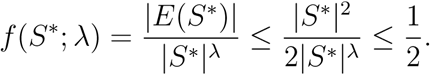

Hence, 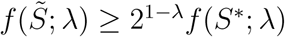 and *c*(*λ*) = 2^*λ−1*^.

**Case 2:** 1 *< λ <* 2. Assume there exists *λ >* 0 such that 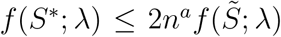 and we want to find such *λ*. It suffices to show

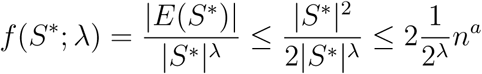

since from case 1, 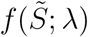 has a lower bound 1/2^*λ*^. It’s automatically true for *|S∗| ≤* (2^2−*λ*^*n*_*a*_)^1/(2*−λ*)^ = 2*n*^*a*/(2*−λ*)^.

For *|S∗*| > 2*n*^*a*/(2*−λ*)^, from the proof of case 2 in lemma 3, we have

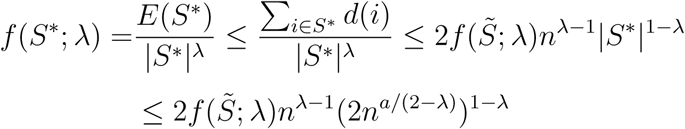

Let 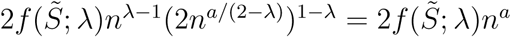, we will get *a* = (*λ −* 1)(2 *− λ*).

Hence, 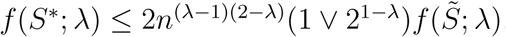, then *c′*(*λ*) = 1 *∨* 21*−λ*.

**Case 3:** For 0.5 *< λ <* 1, the claim is automatically true from lemma 3.

**Case 4:** 0 *< λ <* 0.5

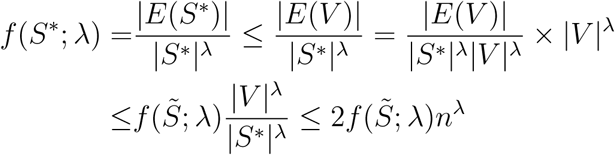

##### 7.3.2 Proof of theorem 2

*Part 1.* Let *𝓒* = {*S*_1_, *S*_2_, *…, S*_*n−*1_} be the sequence of subgraphs generated by deleting the smallest-degree node in Algorithm 1. We first prove the true subgraph *S*_0_ is included in *𝓒* up to a permutation with high probability as *n → ∞*. Denote *n* = *|V |* and *n*_*s*_ = *|S*_0_|.

At stage *k*(*< n − n_s_*) of Algorithm 1, for *i ∈ S*_0_, and *i′ ∈ V/S*_0_, we have

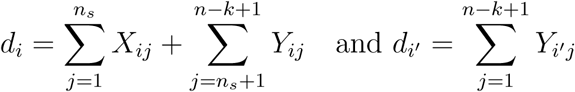

where *X*_*ij*_ ∼ Bernouli(*π*_*s*_), *Y*_*ij*_ ∼ Bernouli(*π*_0_), and *Y*_*i′j*_ ∼ Bernouli(*π*_0_)

From chernoff bound, for *i ∈ S*_0_, *i′ ∈ V/S*_0_ and *δ ∈* (0, 1)

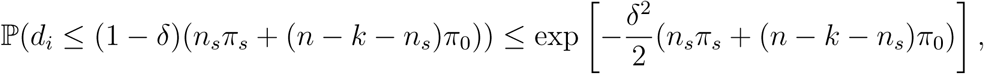

and

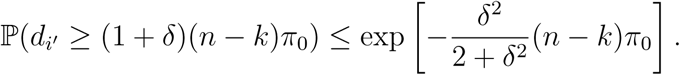

Hence, there exist *ϵ*_0_(*δ*) *>* 0, such that

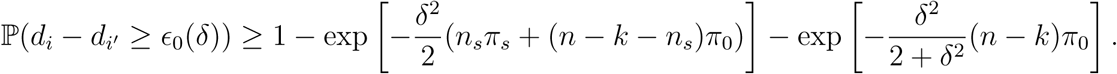

At stage *k*(*< n − n_s_*) of Algorithm 1, the event of deleting a node outside *S*_0_ is:

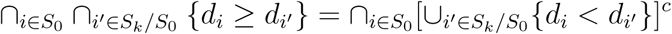

Then,

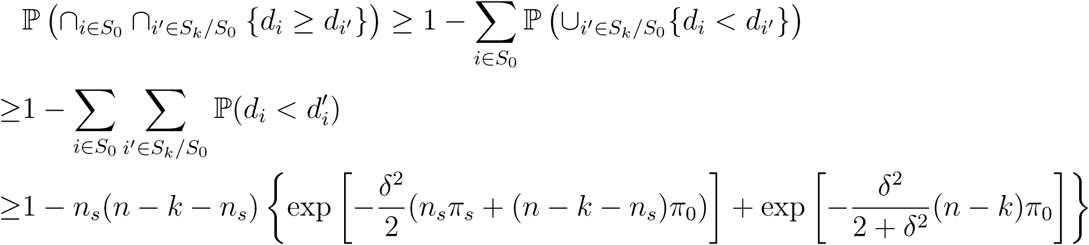

Using similar argument, the class *𝓒* includes the true subgraph up to a permutation *Q*:

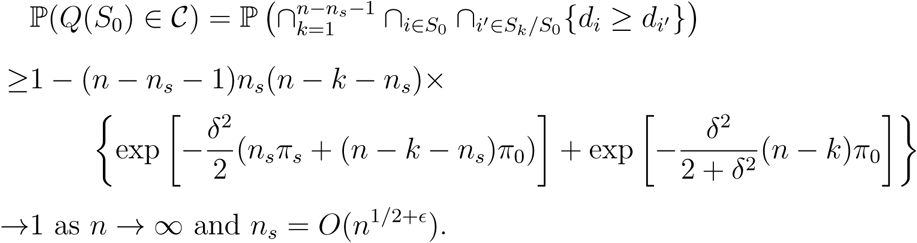

*Part 2.* We then establish the true subgraph can be selected from *𝓒* by density function *f* (*S*; *λ*) for some *λ*. For *k < n − n_s_*, from the proof of part 1, *S*_0_ *⊂ S_k_* with high probability. From Chernouff bounds,

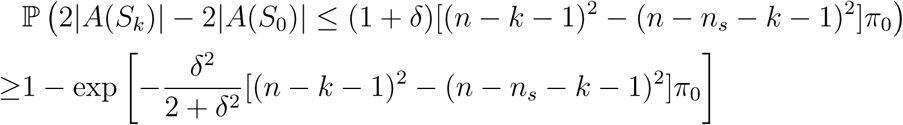

and

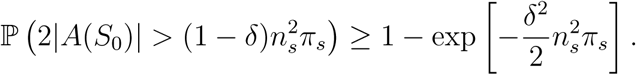

Thus, with high probability,

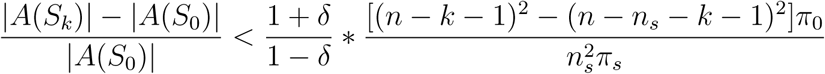

On the other hand,

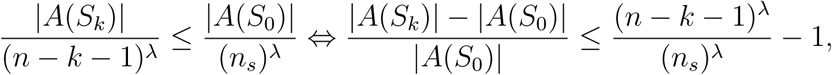

hence, it suffices to have

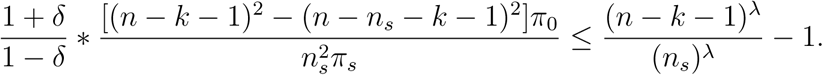

For *π*_*s*_ > *π*_0_, there exist corresponding *δ* and *λ* to make the inequality holds.

For *k* > *n* − *n*_*s*_, we could use similar argument and the claim is true.

##### 7.3.3 Proof of theorem 3

We implement the proof of SBM under maximum likelihood fitting by Choi et al. (2012) that constrain stochastic block model with *K* ≲ *n*^1/2^ communities and the average degree *M* ≳ (log(*n*))^3+*δ*^. The growth restriction on *K* is utilized to bound the number of possible choices of assignment. Although the restriction is not satisfied in our model, the assumption such that all edges outside the dense subgraph are considered as singletons makes the number of possible assignments being the same as *K* = 2. The restriction on average degree is also automatically satisfied for fixed *π*_*s*_ and *π*_0_.

Next, we prove the conclusion is valid when the assignment is maximized over a smaller class of possible solutions, which is generated by different values of *λ*.

The Theorems 1 and 2 in Choi et al. (2012) also hold in our model because the maximization over a subset of parameter space is smaller than over the whole space. For their theorem 3, from the proof of our theorem 2, the true assignment is in the subset that we maximized with probability converging to 1, i.e. 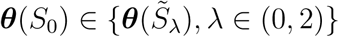. Then theorem 3 holds with probability converging to 1, i.e. 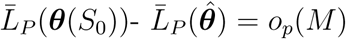. Hence, our claim is true.

##### 7.3.4 Proof of Lemma 1

It’s equivalent to show

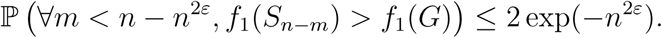

Consider the case *n*_*s*_ = *cn* for some constant 0 *< c <* 1. We first establish *∀m < n − n*^2*ε*^,

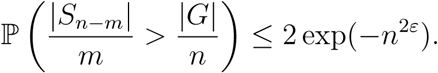

**Case 1:** Consider subgraph *S*_*n−m*_ with *m > n_s_*. We have

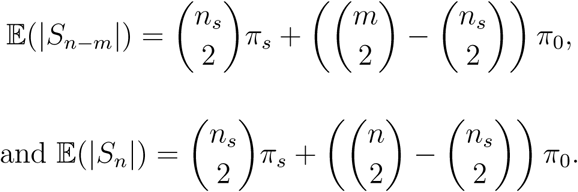

Then, from chernoff bound, we have

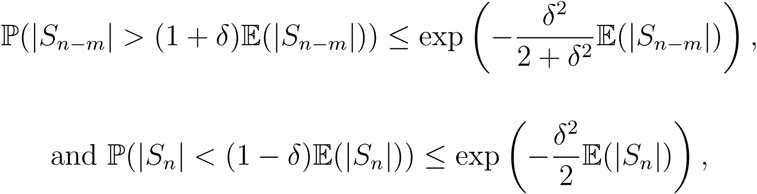

where the second probability is even smaller if we consider the correlation between *|S*_*n−m*_| and *|G|*.

Hence, if we choose *δ* = *n*^*ε*^/*n*, each of the above probability is bounded by exp(*−n*^2*ε*^), in this case, when

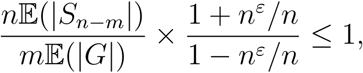

we have

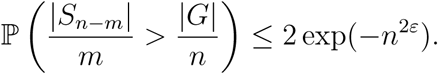

Since 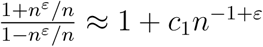, it’s equivalent to have

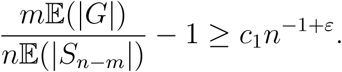

Then,

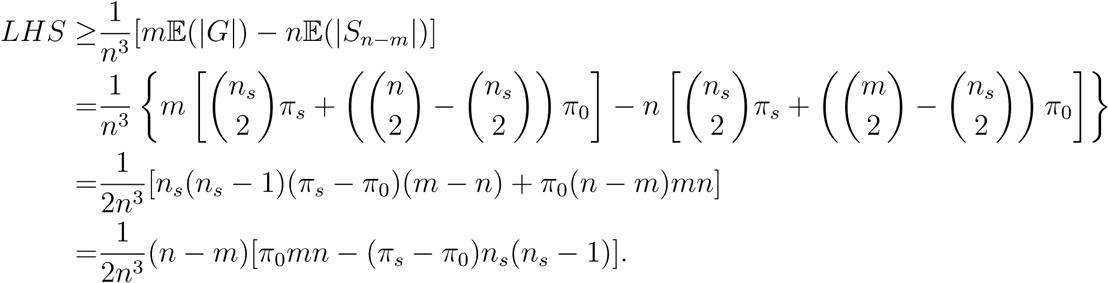

In order to have *LHS ≥ c*_1_*n−*^1+*ε*^, under condition *n − m > n*^2*ε*^, it’s sufficient to have *cπ*_0_ − *c*^2^(*π*_*s*_ − *π*_0_) > 2*c*_1_*n*^*−ε*^

**Case 2:** Consider *m ≤ n_s_*. If

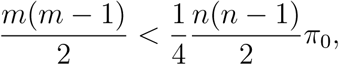

in other words, *m < c*_2_*n* for some *c*_2_, we have 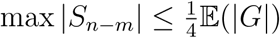, then let *δ* = 3/4,

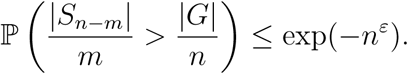

If *c*_2_*n ≤ m ≤ n_s_* = *cn*, using similar argument as Case 1, the claim is true.

The result is true if we take union among all possible *m*.The case with *ns* = *o*(*n*) is automatically true. Hence, the claim is proved.

##### 7.3.5 Proof of Lemma 2

Since *𝓒* = {*S*1, *S*_2_, …., *S*_*n*−2_} is the sequence of subgraphs selected by Greedy Algorithm, for any subgraph in the sequence *S*_*i*_, the next subgraph *S*_*i*−1_ is generated by deleting the node with smallest degree. Without loss of generality, we assume the node deleted from *S*_*i*_ is node *u*. Then, 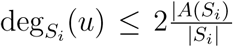, and also, *|A*(*S*_*i*−1_)| = *|A*(*S*_*i*_)| − deg_*S*_*i*__ (*u*). Hence, *|A*(*S*_*i*−1_)| ≥ (1-2/|*S*_*i*_|)*A*(*S*_*i*_)|, and then

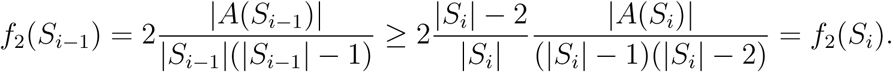

Therefore, the sequence {*S*_1_, *S*_2_, *…., S*_*n−2*_} has nondecreasing values of objective function *f*_2_, i.e. *f*_2_(*S*_*n−2*_) *≤ … ≤ f*_2_(*S*_2_) *≤ f*_2_(*S*_1_). The maximizer of *f*_2_(⋅) has the same value of objective function for all smaller subgraphs in the sequence {*S*_1_, *S*_2_, *…., S*_*n*−2_}.

On the other hand, the smallest subgraph *S*_*n*−2_ can only be two cases: (0, 0; 0, 0) or (0, 1; 1, 0), which has *f*_2_(*S*_*n*−2_) being 0 or 1, respectively. We don’t consider the case with *f*_2_(*S*_*n*−2_) = 0 in general, since from the nondecreasing property, we conclude all subgraphs in the sequence has objective function being 0. For *f*_2_(*S*_*n*−2_) = 1, the nondecreasing property results that the maximizer of *f*_2_(⋅) and all smaller subgraphs have objective function being 1.

Therefore, the densest subgraph must be a clique generated from the graph *G*. Based on the results in Grimmett and McDiarmid (1975), for any random graph *G*(*n, p*), the size of largest clique *K*_*n*_(*p*) in this random graph satisfy,

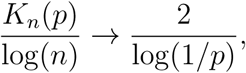

almost surely as *n → ∞*

Since the largest clique generated from the graph *G* with informative subgraph *G*(*S*_0_) is smaller than generated from random graph *G*(*n, π_s_*) with high probability. The densest subgraph returned by Greedy Algorithm in *f*_2_ is bounded by 2 log(*n*)/ log(1*/π_s_*).

## 8 Appendix B: Region Names

### 8.1 Region names

In the following tables, we list the region names and coordinates of subgraph extracted from the data of schizophrenia research.

**Web Table 1:** Region names and coordinates for the nodes in the detected subgraph

| Abbreviation | Region Names | x | y | z |
| --- | --- | --- | --- | --- |
| SFG.R.7.1 | Superior Frontal Gyrus | 7 | 16 | 54 |
| SFG.L.7.5 | Superior Frontal Gyrus | -6 | -5 | 58 |
| SFG.R.7.5 | Superior Frontal Gyrus | 7 | -4 | 60 |
| OrG.R.6.1 | Orbital Gyrus | 6 | 47 | -7 |
| PCL.L.2.1 | Paracentral Lobule | -8 | -38 | 58 |
| PCL.R.2.1 | Paracentral Lobule | 10 | -34 | 54 |
| STG.L.6.2 | Superior Temporal Gyrus | -54 | -32 | 12 |
| STG.R.6.2 | Superior Temporal Gyrus | 54 | -24 | 11 |
| STG.L.6.3 | Superior Temporal Gyrus | -50 | -11 | 1 |
| STG.R.6.3 | Superior Temporal Gyrus | 51 | -4 | -1 |
| STG.R.6.4 | Superior Temporal Gyrus | 66 | -20 | 6 |
| STG.L.6.5 | Superior Temporal Gyrus | -45 | 11 | -20 |
| STG.R.6.5 | Superior Temporal Gyrus | 47 | 12 | -20 |
| STG.L.6.6 | Superior Temporal Gyrus | -55 | -3 | -10 |
| STG.R.6.6 | Superior Temporal Gyrus | 56 | -12 | -5 |
| IPL.L.6.6 | Inferior Parietal Lobule | -53 | -31 | 23 |
| INS.L.6.1 | Insular Gyrus | -36 | -20 | 10 |
| INS.R.6.1 | Insular Gyrus | 37 | -18 | 8 |
| INS.L.6.2 | Insular Gyrus | -32 | 14 | -13 |
| INS.L.6.3 | Insular Gyrus | -34 | 18 | 1 |
| INS.R.6.3 | Insular Gyrus | 36 | 18 | 1 |
| INS.L.6.4 | Insular Gyrus | -38 | -4 | -9 |
| INS.R.6.4 | Insular Gyrus | 39 | -2 | -9 |
| INS.L.6.5 | Insular Gyrus | -38 | -8 | 8 |
| INS.R.6.5 | Insular Gyrus | 39 | -7 | 8 |
| INS.L.6.6 | Insular Gyrus | -38 | 5 | 5 |
| INS.R.6.6 | Insular Gyrus | 38 | 5 | 5 |
| CG.L.7.2 | Cingulate Gyrus | -3 | 8 | 25 |
| CG.R.7.2 | Cingulate Gyrus | 5 | 22 | 12 |
| CG.L.7.3 | Cingulate Gyrus | -6 | 34 | 21 |
| CG.R.7.3 | Cingulate Gyrus | 5 | 28 | 27 |
| CG.L.7.5 | Cingulate Gyrus | -5 | 7 | 37 |
| CG.R.7.5 | Cingulate Gyrus | 4 | 6 | 38 |
| CG.L.7.6 | Cingulate Gyrus | -7 | -23 | 41 |
| CG.R.7.6 | Cingulate Gyrus | 6 | -20 | 40 |
| CG.L.7.7 | Cingulate Gyrus | -4 | 39 | -2 |
| CG.R.7.7 | Cingulate Gyrus | 5 | 41 | 6 |

